# Anterior Insula Reflects Inferential Errors in Value-Based Decision-making and Perception

**DOI:** 10.1101/644427

**Authors:** Leyla Loued-Khenissi, Adrien Pfeuffer, Wolfgang Einhäuser, Kerstin Preuschoff

## Abstract

The process of inference is theorized to underlie neural processes, including value-based decision-making and perception. Value-based decision-making commonly involves deliberation, a time-consuming process that requires conscious consideration of decision variables. Perception, by contrast, is thought to be automatic and effortless. To determine if inference characterizes both perception and value-based decision-making, we directly compared uncertainty signals in visual perception and an economic task using fMRI. We presented the same individuals with different versions of a bi-stable figure (Necker’s cube) and with a gambling task during fMRI acquisition. To track the inferential process assumed in both tasks, we experimentally varied uncertainty, either on perceptual state or financial outcome. We found that inferential errors captured by surprise in the gambling task yielded BOLD responses in the anterior insula, in line with earlier findings. Moreover, we found perceptual risk and surprise in the Necker Cube task yielded similar responses in the anterior insula. These results suggest that uncertainty, irrespective of domain, correlates to a common brain region, the anterior insula. These findings provide empirical evidence that the brain interacts with its environment through inferential processes.

## Introduction

Uncertainty is an unavoidable feature of decision-making. This suggests the brain acts as an inference machine, formulating hypotheses about its environment and testing them against the evidence (Gregory, 1997; Clark 2013). That inference may underlie perception, an unconscious, rapid decision, was posited by the 11^th^ century scientist Alhazen (Howard, 1996) and most famously stated by Hermann von Helmholtz (1867) in the 19^th^ century, who coined the term unconscious inference to describe the process. By contrast, in value-based decision-making, where options are quantified and mulled over by at least one conscious agent, the inferential process includes an explicit, objective component; agents commonly declare their uncertainty in quantitative terms, such as the odds of winning a bet. While several factors differentiate perception from value-based decision-making, they can both be cast as inferential processes, a computation that integrates and resolves for uncertainty. The key question arises: do similar neural mechanisms process uncertainty in value-based decisions and perceptual inference, irrespective of decision features or goals (Parr & Friston, 2017)?

In casting the brain as an inference machine (Friston & Stefan, 2007; Doya et al., 2007), agents are theorized to build internal models that forecast outcomes (Friston, 2010; Clark, 2013). When prediction fails, the error recruits mechanisms to refine the model. Model building and updating is not limited to deliberative judgments, but extends to latent decision-making, as in perception (Haefner et al., 2016). A shift in perception of an unchanging stimulus hints at an internal inferential error (Helmholtz, 1867). As both prediction and error imply uncertainty, decisions cast in a probabilistic framework formally encapsulate an inferential process. Therefore, capturing uncertainty signals in the brain can index a neural inferential process, irrespective of temporal dynamics, consciousness features, or end-goal of a decision.

Perception rarely requires deliberation. Nonetheless, uncertainty arises, as the mapping of the distal world to the sensory signal is not uniquely invertible. Perception infers the world by combining sensory evidence with prior knowledge (Helmholtz, 1867), an approach often cast in Bayesian frameworks (Freeman, 1994; Knill & Pouget, 2004). Perceptual uncertainty has traditionally been captured by drift-diffusion and signal detection models (Heekeren et al., 2008), where reaction time serves as a proxy for decision difficulty, or stimulus uncertainty. In such paradigms, a volitional decision, usually motor, is made when uncertainty is resolved. With multi-stable stimuli however, information conveyed by a constant stimulus is sufficiently uncertain to evoke spontaneous alternations in conscious perception (Rubin, 1921; Boring, 1930; Necker, 1832), suggesting the decision process itself is hidden, with only the outcome made available to the agent. Crucially, ambiguous perception does not rely on dominance time as an index of uncertainty (Brascamp et al., 2005), as switches arise in a stochastic manner., Bi-stable figures have recently been studied in predictive-coding frameworks where perceptual transitions (see also: Dayan, 1998; Hohwy et al., 2008) correlated with anterior insulae and inferior frontal gyrii responses (Weilnhammer et al., 2017), regions also associated with economic uncertainty (Platt & Huettel, 2008; Grinband et al., 2006; Brevers et al, 2015; Preuschoff et al., 2008a; Mohr et al., 2010).

Studies on perception and value-based decision-making have generally remained distinct (Summerfield & Tsetsos, 2012) though some have sought modality-independent neural correlates of the decision process (Ho et al., 2009; Heekeren et al., 2006). Value-based decision-making is driven by utility maximization (von Neumann & Morgenstern, 1945), while successful perception aims for an accurate mental representation of a stimulus. Expected utility violations entail a reward prediction error (Schultz et al., 1997); unexpected shifts in perception should entail perceptual prediction errors (Egner et al., 2010). These domain-specific errors may correlate with distinct neural systems. However, neither utility nor perceptual prediction can elude uncertainty, whose minimization is posited to drive decision-making (Friston et al., 2017). Percept variance is analogous to expected utility risk. Similarly, as risk prediction errors arise in financial paradigms (Preuschoff et al., 2006; Preuschoff et al., 2008a), so they should in perception, when uncertainty breaks through the consciousness barrier, as with bi-stable stimuli.

In previous work, we studied objective expected uncertainty (risk) and its error (surprise) in value-based decision-making with a gambling task, finding a role for the insula in response to both risk and surprise (Preuschoff et al., 2008a). To determine if the same computational account and neural mechanism capture perceptual inference, we sought a task that mimics the probabilistic nature of a gamble but provokes internally generated perceptual errors. To meet this last criterion, we elicited uncertainty with a multi-stable stimulus, the Necker Cube (Necker, 1832), which prompts spontaneous perceptual switches without a corresponding change in stimulus.

Does inference call on a dedicated neural system, irrespective of functional domain? Given the anterior insula’s emergence in perceptual difficulty (Binder et al., 2004; Thielscher & Pessoa, 2007) and in economic uncertainty (Platt & Huettel, 2008; Grinband et al., 2006; Brevers et al, 2015; Preuschoff et al., 2008a; Mohr et al., 2010), we hypothesized this region’s responses correlate with the inferential process. We test this with a common computational account of uncertainty in both perceptual and financial tasks, administered to the same participants during fMRI acquisition. We hypothesized that 1) first order perceptual and reward prediction errors prompt distinct, task-specific BOLD responses in visual and striatal areas, respectively; 2) endogenous (perceptual) and exogenous (economic) inference, as indexed by uncertainty, will correlate with an insular response.

## Materials and Methods

### Recruitment

The study was performed across two sessions scheduled within one week of each other. Sessions were separated due to the considerable length of time spent in the scanner for each (∼45 minutes). The first session comprised the financial uncertainty task (the Card Game, Figure 1a) and the second the perceptual uncertainty task (the Necker Cube Onset task and the Necker Cube Continuous Task) (Figure 1c and Figure 1d). A total of 29 participants were recruited in the Card Game (13 female, 16 male) with a mean age of 25.13 years, of which 4 participants’ datasets were excluded for technical reasons. Twenty-two of the initial pool of 29 participants went on to perform the Necker Cube task. Of these, 19 had also completed the Card Game. In the Necker Cube onset task, 21 subjects were included for analysis (one being excluded for poor behavioral data) and in the Necker Cube continuous task, two further subjects were excluded for inconsistent behavioral data. In total, 18 same participants (8 female, 10 male) completed both sessions to yield usable neuroimaging and behavioral data. The study was approved by the local ethics committee and conformed to the declaration of Helsinki. Participants were recruited through online and paper advertisements broadcast on the Ecole Polytechnique Fédérale de Lausanne and Université de Lausanne campuses. The following inclusion criteria were applied: English speaking; over the age of 18; healthy; normal vision. Exclusion criteria from participation included: history of psychiatric or neurological illness; previous or current psychotropic drug use; metal implants; pregnancy; sensitivity to noise or closed spaces.

**Figure 1.**
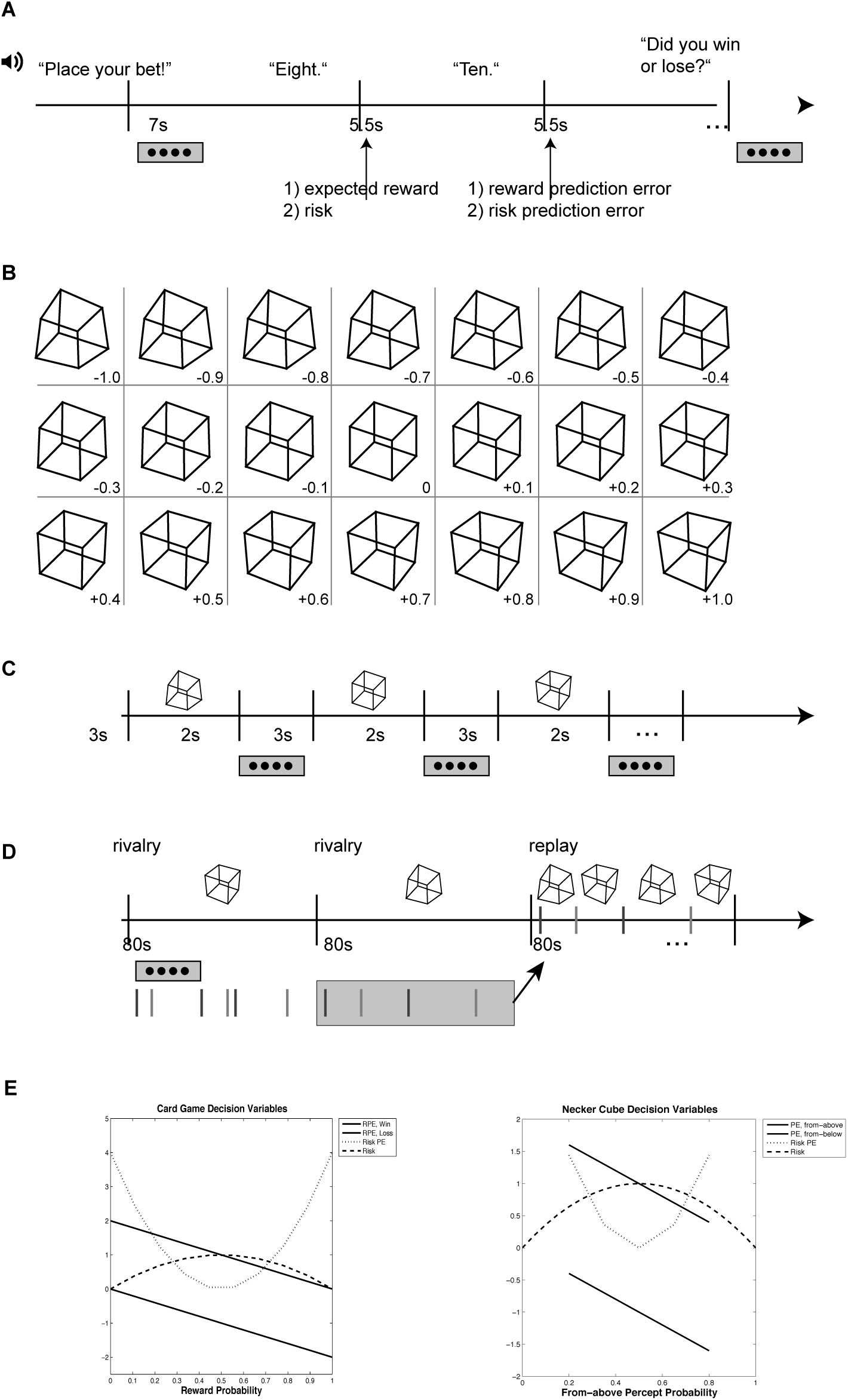
Stimuli and tasks. A) Procedure of the financial uncertainty task (the card game). Participants place their bet (whether the first or the second card will be higher) before the first card is revealed. Decision variables change when cards are drawn, although the decision process is already completed. Participants respond regarding their outcome to ensure they remain vigilant to the task. One trial lasts about 25 s. B) The 21 different versions of the Necker cube used in the onset task, neutral cube in the middle (“0”). C) Procedure of the “onset” Necker cube task. A version of the Necker cube is presented for 3s and participants respond whether they have seen it from above or below in a subsequent 2s interval. D) Procedure of the “continuous” Necker cube task. In “rivalry” phases, participants respond to endogenous changes in the perceptual interpretation of the presented constant Necker cube (adjusted per individual to yield five defined levels of bias in the onset task); in “replay” phases, strongly disambiguated versions mimic the preceding sequence of perceptual switches by exogenous changes. E) Mathematical models of decision variables in the Card Game (left) and Necker Cube (right) Tasks. Both tasks were subject to the same mathematical models of risk and surprise. In the Card Game, risk was computed based on the probability of winning a gamble, given the value of Card 1 (x-axis) and the bet placed, while risk prediction error (surprise) is derived from the difference between the expected risk and actual risk at the outcome of Card 2. The monetary unit on the y-axis is 1 CHF. In the Necker Cube task, risk corresponds to the variance of the probability of viewing a given stimulus from above (x-axis), and surprise computed based on the resulting percept at switch. The two diagrams above show that uncertainty related-variables (risk, surprise) are unsigned and quadratic, whereas first order errors (RPE and perceptual prediction errors) are linear. Most notable is the correspondence of the model across the two tasks to the different paradigms, stimuli and outcomes. (RPE – reward prediction error; Risk PE – risk prediction error; PE perceptual error).

### Procedure

For each of the two sessions, participants were sent digital versions of the study’s information and consent forms on the eve of their study sessions for review. Upon arrival to the study room, participants were queried regarding their understanding of the information before providing their signed consent. Participants were then subject to additional MR safety screening prior to entering the scanning room.

### Tasks

#### Financial Uncertainty Task

We employed a gambling task performed during functional magnetic resonance imaging (fMRI) acquisition. We used an auditory version of a card game (Preuschoff et al., 2006), which has reliably yielded robust results when fit to our chosen computational model. In the task, participants had an equal probability of winning or losing 1 CHF (∼1 USD) at each trial, starting with an initial endowment of 25 CHF. At each round, participants were instructed to place a bet via manual button press on whether a second card drawn from a deck of ten cards would be higher or lower than a first card drawn from the same deck. The bet was made prior to any card being drawn, at which point outcome probability is at 0.5, and the expected value of reward is 0. Following the bet, participants heard the value of the first card. After a 5.5 second interval, participants heard the value of the second card. After another 5.5 s interval, participants were instructed to report whether they had won or lost the round. An incorrect response incurred a penalty of 25 ¢ off a round’s total payoff. Each trial lasted approximately 25 s (Figure 1a). Inter-trial interval durations were randomly jittered (2-5s). All possible card pairs, excluding pairs of identically valued cards, were presented to each participant in a random order, totaling 90 trials per experiment. No two participants played the same sequence of gambles. These 90 trials were divided into three blocks of 30 trials each, to give participants the opportunity to rest between blocks. Each block began with a new 25 CHF endowment. As the task was auditory in nature, once the functional sequence was launched, participants were presented with a black fixation-cross centered on a gray-scale screen throughout the experiment. The task was sounded with the use of Mac OSX text-to-speech function, with the voice of ‘Alex’. Instructions and stimuli were pre-recorded into wav files and transmitted to the participant via MR compatible headphones. The task related session lasted approximately 36 minutes while the imaging session lasted approximately 50 minutes; the whole experimental session, including intake and debriefing lasted approximately 90 minutes.

### Perceptual Uncertainty Tasks

To manipulate perceptual uncertainty, we used different versions of a “Necker Cube” stimulus (figure 1b). Most individuals perceive this wireframe representation of a cube as either a cube seen from-above or from-below, with alternations between the two interpretations. We warped the angles of the cube to bias perceptual interpretations towards one or the other direction.

### Necker Cube, onset task

We presented observers with the 21 versions of the cube depicted in figure 1b. Each version was presented a total of 10 times, in random order (figure 1c). Each presentation lasted 3s, followed by a 2-s blank. Observers were asked to report their percept (“from-above”, “from-below”, as instructed with pictures of extreme versions prior to the experiment) by using two buttons of an MR-compatible response box. To avoid lateralization effects, the assignment of buttons to percepts was counterbalanced across participants. If participants failed to respond within the 2s blank period, the trial was treated as missing data.

### Necker Cube, continuous task

For this task, 5 different versions of the Necker cube were used. These were selected for each individual based on onset-task results. To this end, we fitted a cumulative Gaussian to the dependence of the fraction of “seen from-above” reports on the physical distortion level (cf. figure 2a). Based on this fit, we determined the points at which 20%, 35%, 50%, 65% and 80% views from-above would be reported. (Note that in a few cases, this implied extrapolation to more extreme warpings). Although it cannot be expected that the predominance of the “from-above” view during prolonged viewing (i.e., the percentage of time the “from-above” view is reported dominant) provides a 1-to-1 match to numbers that are based on short presentations, they nonetheless provide a good range in which the respective individual should show variations in pre-dominance around an equi-dominance for both interpretations. For the continuous task, we therefore presented those 5 stimulus versions to participants that corresponded to their individual 20%, 35%, 50%, 65%and 80% fits from the onset task. These stimuli were presented in 11 trials, for 80 seconds at a time (figure 1d). The 50% stimulus was shown in the first, the sixth and the eleventh trial, the order of the other stimuli was random (one presentation each in trials 2 through 5, one in trials 7 through 10). Participants were asked to report switches of their perception by a button press, one button for switches towards the “from-below”, another for switches towards the “from-above” state. Crucially, participants were asked to report switches when they arose, if they arose, to encourage a passive viewing and experience of perceptual switches. Button assignment in each individual was consistent with the onset task. Four of the 11 trials (one per cube version, randomly picked), were followed by a replay phase, in which strongly disambiguated Necker Cube versions were presented to induce perceptual changes exogenously at the same time-points as reports in the previous experimental phase. Replay phases were excluded from the analysis of behavioral data, as were rare occasions of reported switches that were followed by a “switch” to the same state (i.e., the same button was pressed twice in succession). The latter were also removed from imaging analysis (see below).

**Figure 2.**
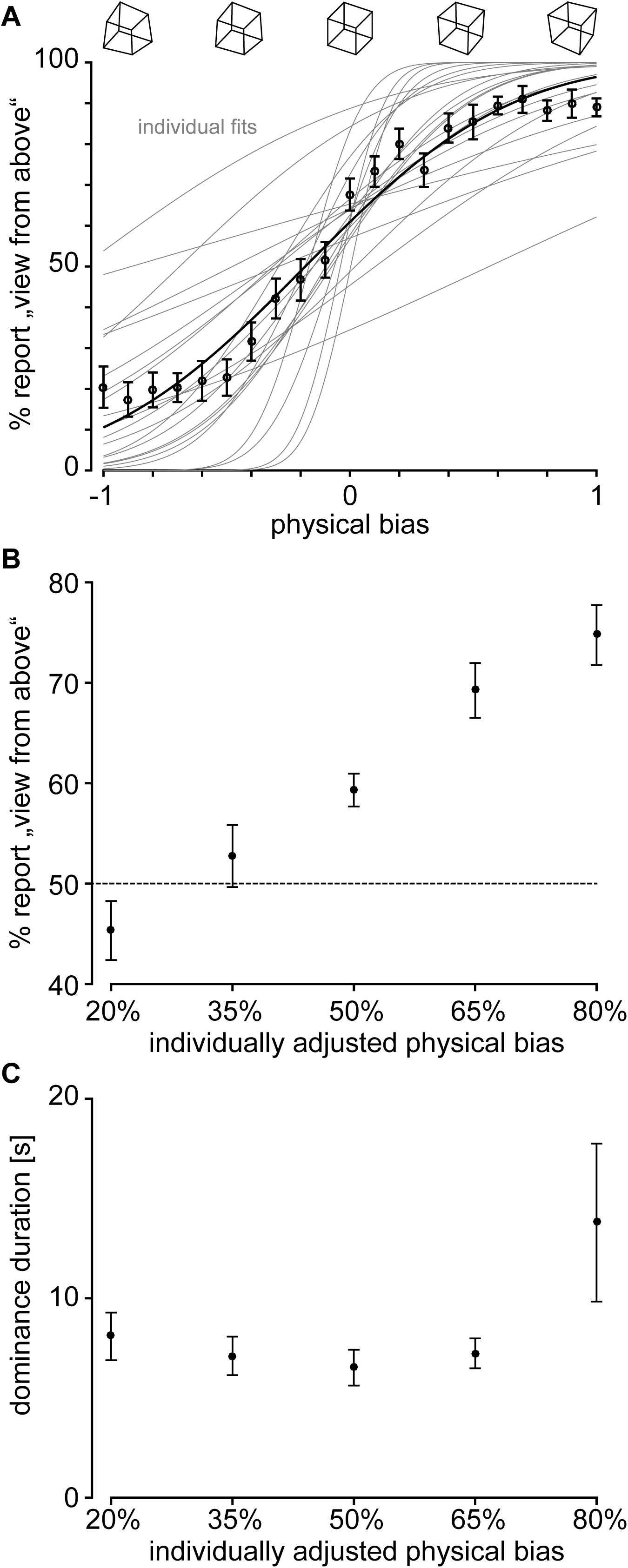
Behavioral data of the Necker cube tasks. A) Onset task: Fraction of “from above” reports for each of the 21 cubes (figure 1c) were fitted by a cumulative Gaussian per individual (gray lines), and to the average (black symbols, mean +/- standard error of mean (sem) across subjects; black line: fit to aggregated data. B) Continuous task: predominance of the ‘from above’ percept (i.e., time the ‘from above’ precept was reported divided by the sum of times either ‘from above’ or ‘from below’ was reported). x-axis denotes individual adjusted ‘from above’ level according to onset task. Mean and sem across participants. C) Median duration of a dominance phase, mean and sem across observers, x-axis as in panel B.

All tasks were coded in Matlab (Matlab and Statistics Toolbox Release 2013a, TheMathWorks, Inc., Natick, Massachusetts, United States) and the Psychophysics Toolbox (Brainard, 1997; Pelli, 1997; Kleiner et al, 2007).

### Image acquisition

Scans were acquired on a Siemens 3T Prisma at the Centre Hospitalier Universitaire Vaudois. Once settled in the bore, we acquired first a localizer scan, followed by a gre-field mapping scan to generate voxel displacement maps (64 slices; 3 × 3 × 2.5 mm resolution; FOV 192 mm; slice TR/TE 700/4.92 ms; FA = 80 degrees; Base resolution = 64 mm); and a B1 mapping scan to correct for magnetic field inhomogeneities (4 × 4 × 4 mm resolution; FOV = 256 mm; slice TR/TE 200/39.1 ms; Base resolution 64 mm). We then alerted participants to the beginning of the task and functional acquisition. Parameters for our EPI sequence were as follows: 2D EPI, Multi-Echo sequence (3 echo times), 3 × 3 × 2.5 mm resolution, FOV = 192 mm; FA = 90 degrees, slice TR = 80 ms; TE = (17.4; 35.2; 53 ms); base resolution 64 mm. At the end of the experimental session, anatomical T1 images were acquired with the following sequence parameters: T1 MPRAGE, 1×1×1 mm resolution; FOV = 256 mm; slice TR/TE = 2 ms/2.39 ms; FA = 9 degrees; base resolution = 256 mm).

### Image preprocessing

Functional scans were preprocessed and analyzed using SPM 12. As we employed a multi-echo EPI sequence with 3 echo times, we first summed the three volumes to obtain one scan per TR (Gowland & Bowtell, 2007). We applied slice-timing correction to the volumes, as they were acquired with a comparatively long effective TR (2.72 s) (Sladky et al., 2011). We then generated voxel displacement maps (VDM) and applied these to functional volumes. Volumes were warped and realigned to the mean functional image before bias-field correction. Then data were co-registered to individual T1-weighted anatomical volumes before being segmented (6 class tissue probability maps) and normalized to the Montreal Neurological Institute 152 template. Volumes were then smoothed with a Gaussian kernel of 6 mm FWHM (Friston et al., 1995). The resulting images were used for analysis.

### Mathematical Models of Risk and Surprise

Mathematical models of risk and surprise for the Card Game have been described elsewhere (Preuschoff et al., 2006; Preuschoff et al., 2008a; Preuschoff et al., 2011). Our mathematical model of risk is analogous to mathematical models of entropy (Quartz, 2009; Faraji et al., 2018). We use risk prediction error as a measure of surprise, which yields values that are highly correlated with alternative models of surprise, notably absolute prediction error (Preuschoff, et al., 2011) and Shannon information (Strange et al., 2005) (see Appendix A.4). Therefore we do not consider alternative accounts of risk or surprise as competing models to our study. We borrow the same accounts used in our previous studies and adapt them to our perceptual uncertainty task in order to build on previous results relating to economic decision-making (Figure 1e). Our measures of perceptual uncertainty are based on the variance of the probability distribution associated with viewing the cube from-above. Each Necker Cube stimulus is endowed with an expected value, the average of the probability of viewing a cube from-above and the probability of viewing the cube from-below. We derive 2 measures of perceptual risk associated with a Necker Cube stimulus, one objective and one subjective. For subjective risk, we compute the squared difference of a probability associated with viewing a given stimulus from-above from the expected value of that same stimulus squared, based on individual responses in the Necker Cube – onset task. Objective risk is derived by computing the same formula, but with assumed probabilities of viewing the cube from above based on the cube’s warp bias. These measures mirror risk accounts used in economic contexts but for one difference; expected values in the latter are associated with a monetary gain, or reward. Here, we substitute a monetary gain value of +1 with a from-above perceptual outcome of +1; similarly, we substitute a monetary loss value of -1 with a from-below perceptual outcome of -1. Finally, we derive a prediction error related to the switch by subtracting the stimulus’ expected value from the perceptual state following the perceptual switch; and a risk prediction error captured by the difference between the squared prediction error and the stimulus-associated risk value.

**Equation 1: Predicted Reward**

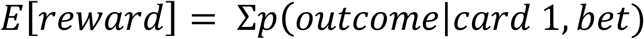

**Equation 2: Reward Prediction Error**

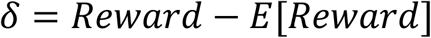

**Equation 3: Risk Prediction Error (Surprise)**

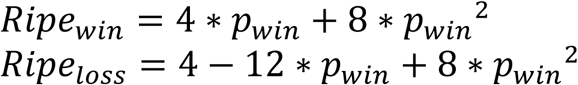

**Equation 4 Expected Percept**

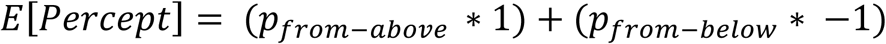

**Equation 5 Perceptual Risk**

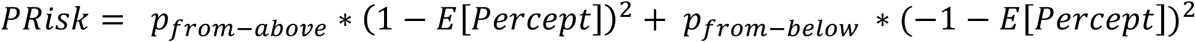

**Equation 6 Surprise As Risk Prediction Error for Perceptual Switches**

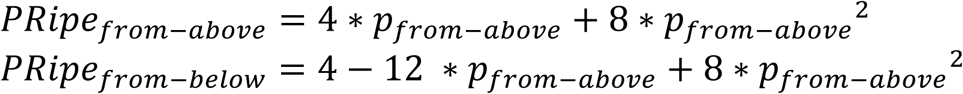

### fMRI Data analysis

Functional data analysis was implemented in SPM12b (www.fil.ion.ucl.ac.uk/spm). We performed a model-based fMRI analysis on all 3 experimental tasks (Card game, Necker onset, Necker continuous) using a summary statistic approach (Worsley et al., 2002). To address our research question, we first analyzed individual subject EPI time-series using a fixed-effects general linear model (GLM) before pooling subject-level contrast images in a random effects analysis, where contrasts of interest were examined with one-sample t-tests to obtain an average response. Regressors in the respective GLMs were convolved with the hemodynamic response function and parametrically modulated by decision variables. We applied the same computational models to both the financial and perceptual uncertainty paradigms via parametric modulation. Each parametric modulator was automatically orthogonalized with respect to the event onset and sequentially to prior parametric modulators. Below, we describe GLMs applied to each task in greater detail.

### Financial Uncertainty

In the card game, we modeled the 5.5 s period after the participant hears the value of card 1 as a boxcar function and further parametrically modulated these events with expected reward followed by a risk value; and we further modeled the 5.5 s period after the participants hears the value of card 2 with a boxcar function and parametrically modulated these epochs with a reward prediction error followed by a risk prediction error. The model also included two nuisance regressors including events modeled as delta functions, including one for button press events (0s duration), to report the bet and the trial outcome, and one for sound, upon hearing instructions for the bet placement and the trial report (0s) duration (Figure 3c). These regressors were convolved with the canonical hemodynamic response function (HRF). Expected reward and risk, as well as reward prediction error and risk prediction error are not expected to correlate (Figure 1e) and show little evidence of multicollinearity in subject-level design matrices. These quantities are further subject to orthogonalization in the SPM software. Also included in the GLM were 6 movement related regressors of no interest.

**Figure 3.**
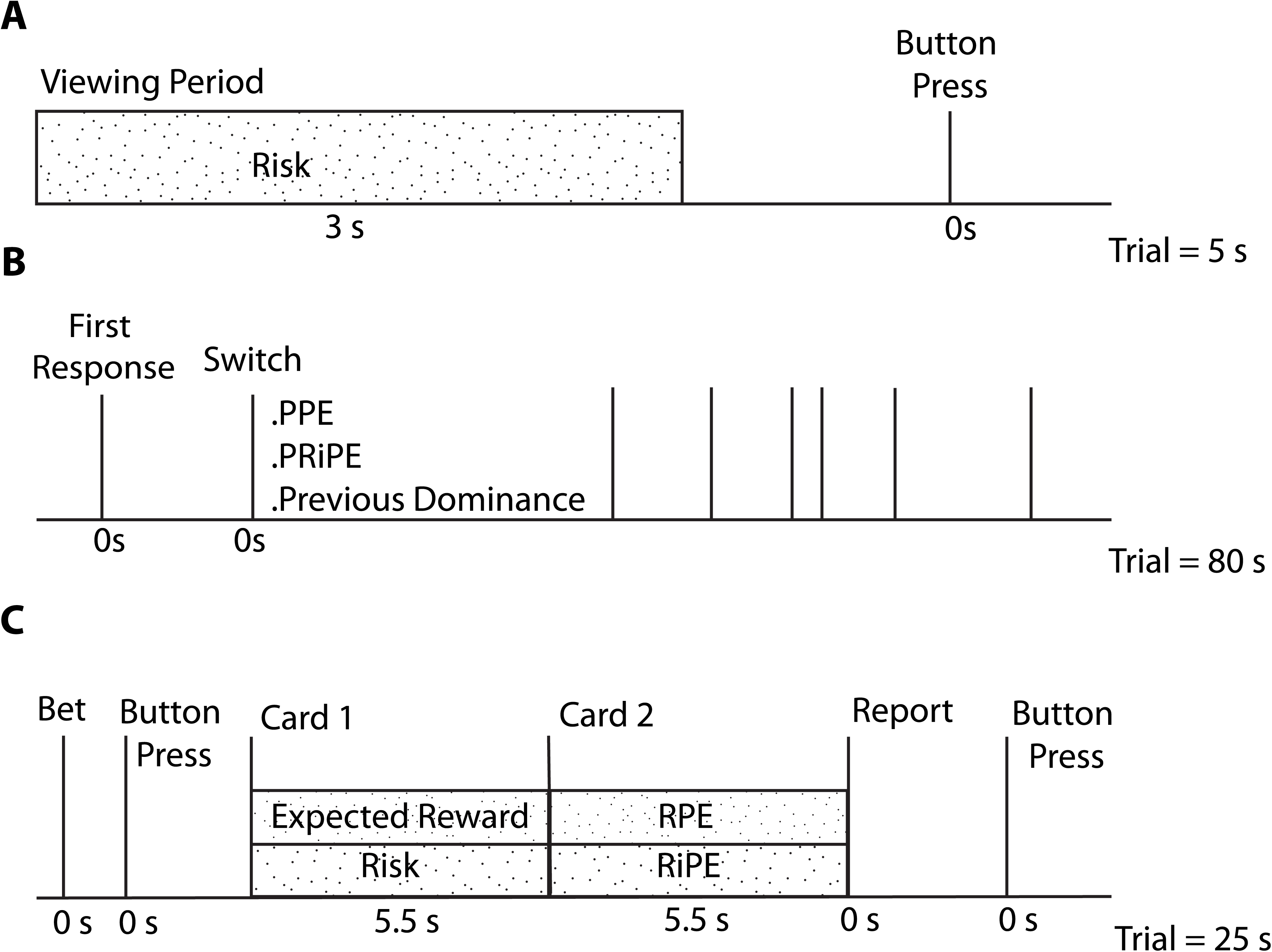
Graphical representation of general linear models for the three tasks. A) The Necker Cube Onset task GLM was comprised of a regressor for the viewing period of the cube (3s) parametrically modulated by stimulus risk, as well as a regressor including button responses. B) The Necker Cube Continuous Task included regressors for first response onsets, and a regressor for switch responses, parametrically modulated by first order perceptual prediction error; perceptual risk prediction error; and previous dominance time. C) The Card Game GLM included two regressors of no interest comprised of button presses (for bet and trial outcome), modeled as stick functions; and of sound (for request to bet and to report trial outcome), also modeled as stick functions. The sounding of cards 1 and 2 were modeled with boxcar functions (5.5s durations), parametrically modulated by expected reward and risk; and Reward Prediction and Risk Prediction Error, respectively. (PPE: perceptual prediction error: PRiPE: perceptual risk prediction error; RPE: reward prediction error; RiPE: risk prediction error)

### Necker Cube - onset task

We constructed 2 general linear models containing 2 experimental regressors as well as 6 movement-related regressors of no interest. The GLMs differed along one dimension; specifically, in one GLM, perceptual risk was computed based on objective probabilities of viewing a trial-related stimulus from-above, or probabilities related to the degree of experimentally manipulated cube bias towards one percept or another; while in the other, perceptual risk values were derived from subjective probabilities of perceiving one cube representation or another, computed at the end of the task from individual participants’ reports. The first regressor in both GLMs contained onsets for stimulus presentation, modeled as boxcar functions of 3 seconds (the viewing period), parametrically modulated by risk values. The second regressor in the GLM contained onsets for button presses during the response window modeled as delta functions. Regressors above were convolved with the canonical HRF. In addition, the model included six movement-related regressors of no interest (Figure 3a).

### Perceptual Surprise, Necker Cube continuous task

In the continuous task of the perceptual uncertainty experiment, we constructed a GLM with the following regressors. Our primary regressor of interest was composed of perceptual switches modeled as delta functions, parametrically modulated by the perceptual prediction error, followed by the risk prediction error, or surprise, and finally by the amount of time spent in the previous perceptual state, to control for potential time-related effects. The risk prediction error was computed based on the participant’s individual probability of seeing a stimulus from-above or from-below and modeled each switch by this value given both the stimulus and the perceptual state into which the participants switched to. The 5 different levels of cube warping towards a from-above state (20%, 35%, 50%, 65% and 80%) yielded 5 values of surprise, with maximal surprise emerging for least likely perceptual states in most disambiguated stimuli (e.g., switching into a “from-below” perceptual state for a cube biased at 80%). We further included replay condition presses as a regressor in the model to account for BOLD responses to exogenous stimulus switches. These regressors were convolved with the canonical HRF. We also included 6 motion-related regressors of no interest in the GLM (Figure 3b).

## Results

### Financial Uncertainty Task - Behavioral Results

Of 29 participants, four were excluded for technical reasons. Twenty-five remaining participants were included in the analysis of the financial uncertainty task. Participants were paid for one out of the three blocks performed. Average task-related payout per participant was 29.57 CHF; across all blocks and participants, payoffs were in the range of 13-39 CHF.

### Perceptual Uncertainty Tasks - Behavioral Results

In the onset task, participants were briefly presented 21 different versions of the Necker Cube and the percentage of “view from-above” was determined across 10 presentations of each. One participant, who classified all versions of the Necker as seen from-above, was excluded from further analysis. As intended, “from-above” percepts increased with warping level and were well fitted by a cumulative Gaussian function (Figure 2a, gray). The “point of subjective equality” – i.e., the warping at which 50% “from-above” was perceived - was slightly shifted towards warpings indicating “from-below” (Figure 2a). This is consistent with a “from-above” bias for the symmetric configuration.

From the individual fits, we determined the cube versions at which the individual had 20%, 35%, 50%, 65% and 80% dominance for the “view from-above” at onset. The corresponding cubes were used for the continuous task in the respective individual. From the analysis of this task, one additional participant who on average reported less than 1 perceptual transition per condition, was excluded. Since the task differed between the onset task (respond to a cube onset) and the continuous task (report switches during continuous viewing) it could not be expected that predominance of the from-above percept (i.e., the percentage of time the from-above percept is seen as dominant) would match 1-to-1 to the nominal values of 20%…80%. Nonetheless, we found that the manipulation was successful: in the rivalry conditions of the continuous task, there was a significant main effect of manipulation on predominance (F(4,80)=22.8, p<.001; repeated-measures ANOVA) and the increase with the manipulation level was monotonic and as intended (Figure 2b). Importantly, there was also a significant main effect of the cube manipulation on dominance durations (F(4,80)=3.53, p=.01) with a minimum at the intermediate manipulation (“50%”) and increasing values towards the more extreme cubes (Figure 2c). This is consistent with the intended experimental manipulation: high levels of uncertainty at the “50%” level, medium levels of uncertainty at the “35%” and “65%” level and low levels of uncertainty at the “20%” and “80%” level. We fit dominance times from all individuals to a gamma distribution to determine the temporal characteristics of perceptual switches (Zhou et al., 2004) and find that Necker Cubes across all bias values conform to such a distribution (Figure A.3).

### Imaging Results

Functional data analysis was implemented in SPM12b (www.fil.ion.ucl.ac.uk/spm) at the single subject level, and the SnPM toolbox (http://warwick.ac.uk/snpm) for group level analyses. Our second level analysis comprised whole-brain permutation tests (10 000 permutations), with a variance smoothing of 6 mm. We applied a false discovery rate (FDR; Benjamini & Hochberg, 1995) threshold of 0.05 at the voxel-level to control for false discovery rates (Hahn & Glenn, 2018). For completeness, we also report results using a whole-brain family-wise error corrected threshold of p = 0.05 (Eklund et al., 2016).

### Financial Uncertainty

#### Contrast Reward Prediction Error (RPE)

We performed a one-sample t-test on the parametrically modulated outcome of each trial by its corresponding RPE value. A positive signed t-test on this regressor using FDR correction yields significant clusters in expected areas, notably bilaterally in the caudate (right and left, p = 0.009 and 0.015, t =4.26 and 3.95, k = 250 and 361) and the cingulate cortex (left middle and right posterior, p =0.017 and 0.022, t =3.93 and 3.87, k = 74 and 67). We further find significant clusters bilaterally in the inferior frontal gyrus (p =0.009 and 0.009, t =6.94 and 4.86, k = 7910 and 1012), left hippocampus (p = 0.009, k = 75, t =3.72); and right middle temporal gyrus (p = 0.011, t =4.21 and k = 114) (Figure 4a). When controlling for FWE, we find significant peaks in right inferior frontal gyrus (p < 0.001; t = 6.94; k = 150); left medial frontal cortex (p =0.004; t = 5.57; k = 202); left superior frontal gyrus (p = 0.022; t = 4.92; k = 10); left middle temporal gyrus (p = 0.036; t = 4.74; k = 5).

**Figure 4.**
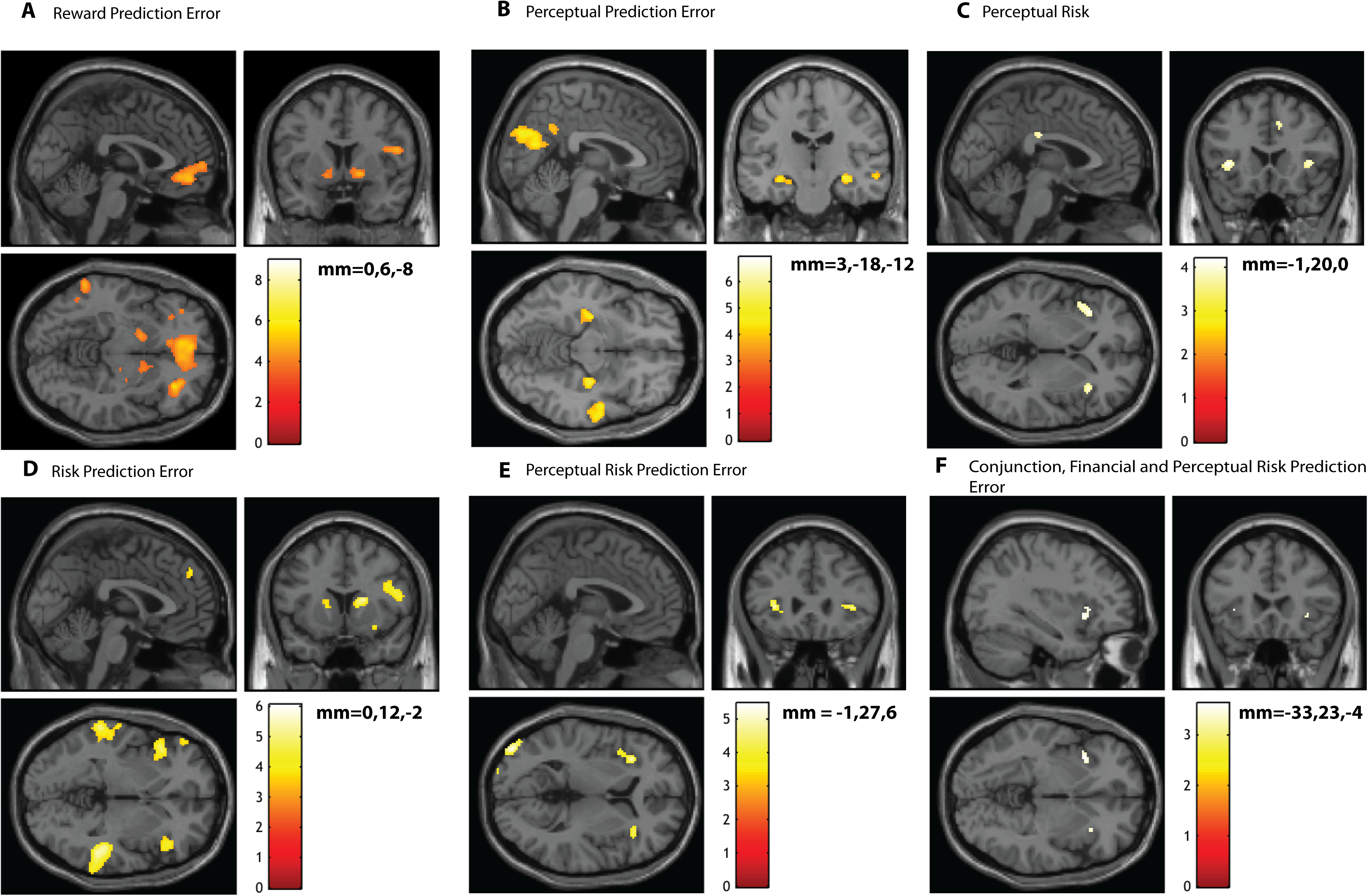
A) Contrast for reward prediction error in the Card Game at the gamble’s outcome shows an expected pattern of BOLD responses in the dorsal striatum (bilateral caudate) and cingulate. B) Perceptual prediction errors in the Necker Cube Continuous task show significant clusters in bilateral. hippocampus, and the cuneus and also yielded several significant clusters in visual areas. C) fMRI results for Perceptual Risk in Necker Cube Onset Task. Risk modulation in perceptual uncertainty yields significant clusters in the bilateral insula. The viewing period of the Necker Cube (3 s) was modulated by a measure of subjective risk, based on the probability of a participant viewing a particular Necker Cube stimulus from above or from below. Images show non-parametric statistical maps thresholded at p =0.05, FDR corrected. D) Surprise contrasts in the financial uncertainty paradigm find significant BOLD responses in the bilateral anterior insula, and middle temporal gyrii and right caudate, replicating previous findings. Shown are non-parametric statistical maps thresholded at p =0.05, FDR corrected. E) The contrast for surprise modeled at the perceptual switch response in the Necker Cube Continuous task yields significant BOLD responses in the anterior insula as well as in occipital regions. Shown are non-parametric statistical maps thresholded at p =0.05, FDR corrected. F) Thresholded maps are shown for the conjunction analyses of surprise contrasts in the financial and perceptual tasks. These contrasts were tested against the conjunction null and the analysis constrained to one anatomical map of both left and right anterior insulae. All colorbars above represent t-values.

#### Contrast Risk Prediction Error (RiPE)

Trial outcomes at Card 2 were further modulated by their risk prediction errors (surprise) in addition to their reward prediction errors. We replicate previously reported results for surprise at Card 2, where we find significant clusters in the left and right insulae (p =0.031 and 0.031, t =4.70 and 4.56, k = 870 and 1597) and the right caudate (p =0.029, t =4.96, k = 293), as well as left and right medial temporal gyrii (p =0.029 and 0.029, t =6.11 and 5.33, k = 1237 and 694) (Figure 4d). While memory regions are not of direct interest to our research question, significant clusters in the medial temporal gyrii support the notion that surprise acts as a learning signal. Controlling the FWE rate, we find significant responses in left and right middle temporal gyrus (p < 0.001 and p = 0.006; t = 6.11 and t = 5.33; k = 118 and k = 32); and right caudate (p = 0.019; t = 4.96; k = 6).

### Perceptual Uncertainty

#### Onset task – contrast risk

We performed a t-test on the viewing period of the cube (3s) prior to the participant’s percept selection, modulated by the stimulus risk to probe perceptual risk activation. Perceptual risk was defined as the variance related to the average perceptual value of a stimulus, based on the probabilities of its being viewed from-above or from-below. Using experimentally derived subjective risk values, whole brain analysis reveals responses in the left and right anterior insula (p =0.009, t =4.10, k = 74; p= 0.014, t =3.95, k =44) (Figure 4c). Objective risk values yielded no significant voxels, underlining the importance of accounting for individual differences in perception. Using a FWE error rate on the same data, we find the same significant clusters in left and right insula (p = 0.009 and p =0.014; t = 4.10 and t = 3.95; k = 74 and k = 44).

#### Continuous task - Contrast Perceptual Prediction error

In the Necker Cube Continuous task, we modulated the manual response to a perceptual switch with 3 values: the first-order perceptual prediction error; the time spent in the previous perceptual state, to account for BOLD responses related to dominance times; and a perceptual risk prediction error. We first performed a contrast on the perceptual prediction error, to account for a first-order error as we do in the Card Game for reward prediction error. Results show significant clusters in the precuneus (p = 0.035, t =5.20, k = 2231), left and right angular gyrii (p = 0.035 and 0. 046, t =4.75 and 4.34, k = 882 and 19), left and right occipital poles (p = 0.040 and 0.035, t =4.24 and 4.41, k =32 and 68), left and right hippocampi (p = 0.035 and 0.035, t =4.34 and 4.56, k = 232 and 122), and right middle temporal gyrii (p = 0.035, t =4.75, k =392) for this contrast (Figure 4b). Thus the pattern here shows a response in memory regions (hippocampus and middle temporal gyrus) in addition to the visual system, suggesting the error calls on higher-level functions such as learning, in addition to perceptual processes. Applying a FWE correction yields significant results in the left precuneus (p = 0.024; t = 5.20; k = 31).

#### Continuous task - Contrast Perceptual Risk Prediction error (Surprise)

A positive contrast for the perceptual risk prediction error yielded significant clusters in several regions of the visual system, including bilateral mid, inferior and fusiform occipital gyrii and left calcarine cortex. In addition however, significant BOLD responses were found bilaterally in the anterior insulae (p = 0.035 and 0.040, t =4.34 and 3.85, k = 172 and 56) (Figure 4e). Using FWE, we find significant responses in the left middle occipital gyrus (p = 0.014; t = 5.34; k = 38); right precentral gyrus (p = 0.025; t = 5.07; k = 8); right occipital fusiform gyrus (p = 0.029; t = 5.01; k = 12); and left inferior occipital gyrus (p = 0.030; t = 4.99; k = 8).

#### Financial and Perceptual Risk Prediction Error (Surprise) – Conjunction Analysis

To test for a common BOLD response to surprise across the two tasks, we performed a conjunction analysis of the surprise contrasts in the financial and perceptual paradigms (N = 25 and N = 21, respectively). We performed this analysis in SPM by selecting a two-sample t-test design for non-independent samples of unequal variance. Though both our initial hypothesis and subsequent results highlight the insula as a prime region of interest for our research question, we opted to perform this test at the whole brain-level in a first instance. We tested against the conjunction null (logical AND) (Friston et al., 2005) and found a significant cluster in the left anterior insula (p = 0.042, FWE corrected; k = 33; peak voxel coordinates -36, 16, -4) as well as a non-significant cluster in the right anterior insula (p = 0.078, FWE corrected; k = 8; peak voxel coordinates (34, 24, -4) (Figure 4f). Crucially, no other voxels or clusters emerged in the conjunction analysis, highlighting the shared representation for surprise across the tasks in the insula only.

### Direct Within-Participant Comparison of Insular BOLD Responses to Surprise Across Tasks

Neuroimaging results above show first that an account of perceptual risk yields a BOLD response in the insula, as reported in previous fMRI studies investigating financial risk. Further, we find insular responses for surprise in both our perceptual and financial tasks. As our experimental paradigm is a within-participant design, we next compared surprise contrast beta values in the anterior insula for participants who performed both experiments and whose data were not excluded from analysis (N =18) to directly compare BOLD responses for surprise across the two tasks, in spite of inter-task and inter-session effects (Gonzalez-Castillo et al., 2017). We performed this analysis by extracting individual beta values from the peak voxels reported in the conjunction analysis above, for left and right anterior insula, as well as the average beta value for each individual in the cerebellar vermis (VI-VII lobules) using anatomical masks from the Neuromorphometrics Atlas (Neuromorphometrics, Inc.) and the MarsBar toolbox (Brett et al., 2002). We selected the cerebrellum as a reference region because it is a midline region that is likely not involved in higher-level decision-making processes. Individual beta values for each of these three regions were plotted against each other across the two tasks for surprise to determine if BOLD responses for surprise co-varied in the same direction. We find a positive, non significant relationship in left and right anterior insula across the two tasks (r(16) = 0.33, p =0.18 for left insula; r(16)= 0.26, p =0.30 for right insula; r(16) =0.44, p =0.065 for left insula in financial surprise against right insula in perceptual surprise and r(16) = 0.17, p = 0.50 for right insula in financial surprise against left insula in perceptual surprise) (Figure 5a). By contrast, individual average beta values for surprise in the reference region show no relationship across tasks (r(16) = -0.05, p = 0.84) (Figure 5b).

**Figure 5.**
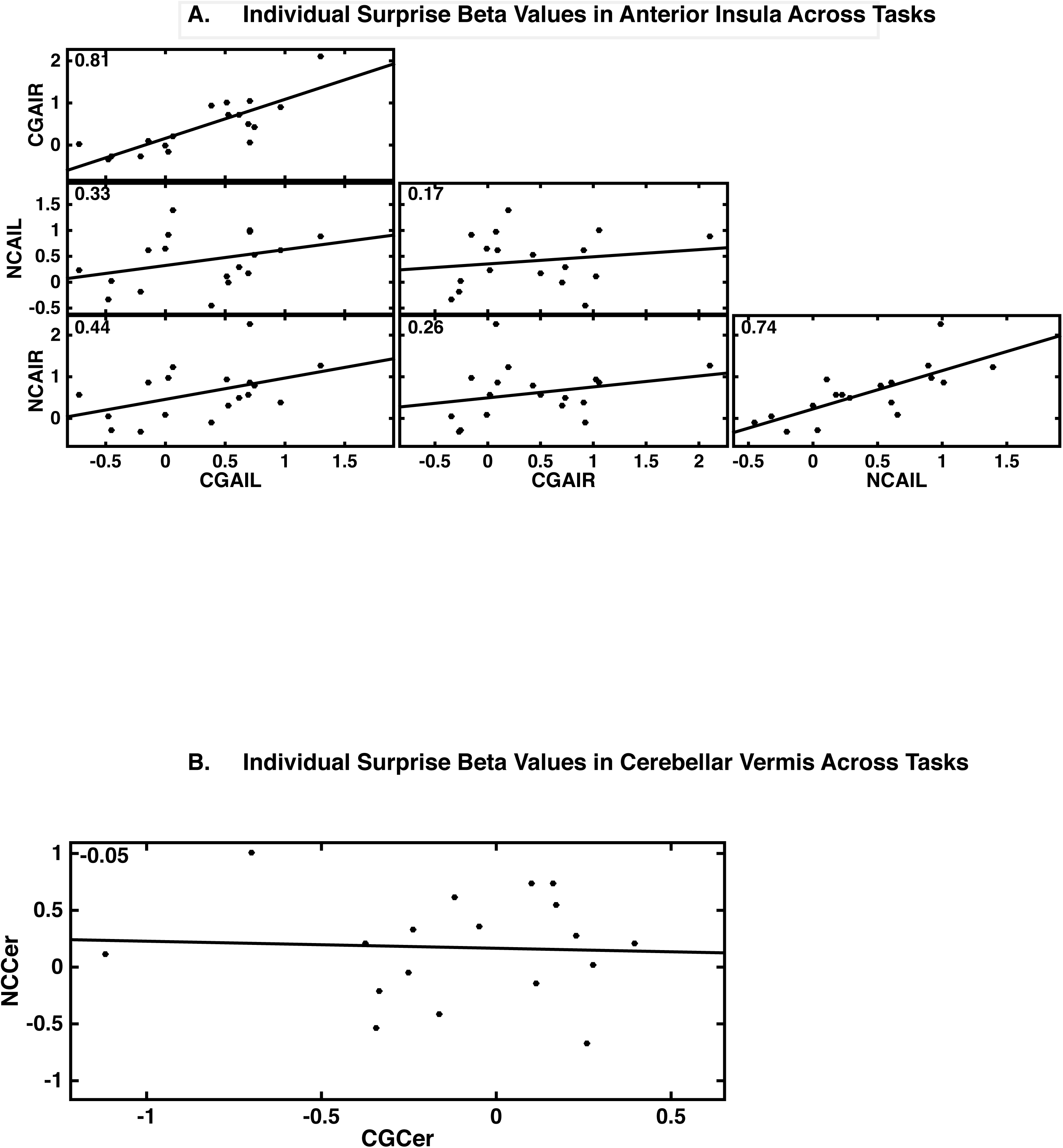
A) Individual subject’s average beta values for surprise regressors in perceptual and financial uncertainty were extracted in each of the following regions: left and right insulae. We directly compared the anterior insula’s involvement in surprise in a within-subject correlation (N=18). Results show a non-significant positive correlation across tasks in the anterior insula. B) Individual subject’s average beta values for surprise contrasts in perceptual and financial uncertainty within the cerebellar vermis (VI-VIII) were plotted. In contrast to the correlation plot of the insula, we find no correlation across tasks for surprise in the cerebellum vermis (CGAIL – Card Game Anterior Insula Left; CGAIR – Card Game Anterior Insula Right; NCAIL – Necker Cube Anterior Insula Left; NCAIR – Necker Cube Anterior Insula Right; CGCer – Card Game Cerebellum; NCCer – Necker Cube Cerebellum).

## Discussion

To probe the presence of a generalized neural correlate of the inferential process in both value-based decision-making and perception, we applied a formal account of economic risk (Preuschoff et al., 2008b) to BOLD signals arising from uncertainty in financial and perceptual domains. Using the Necker Cube to elicit perceptual uncertainty, behavioral results show that experimental manipulation of the stimulus succeeded in expanding the range of probabilities associated with two percepts, allowing for a derivation of “perceptual risk”. To capture the endogenous inferential process related to perception, we model fMRI BOLD responses to perceptual switches with an economic account of uncertainty, finding a significant response in the anterior insula to perceptual risk and its error, surprise. We further administered a well-established gambling task in the same individuals, replicating results in the anterior insula for surprise. A whole-brain conjunction analysis of the two tasks finds significant clusters exclusively in the anterior insula. Finally, we found a positive relationship between individual insular BOLD responses and surprise across tasks. Our results suggest a common neural system dedicated to processing uncertainty in the anterior insula and more generally, support the theoretical framework of the brain as an inference machine (Friston, 2010), irrespective of stimulus or goal features.

Our tasks differed in several dimensions, retaining a commonality in uncertainty and hence, an unavoidable inferential process. Results are therefore remarkable because we lift an objective, economic model of uncertainty and apply it to a perceptual paradigm, with all that the latter implies: spontaneity, subjectivity and absence of conscious deliberation. In the Card Game, surprise arises in response to a changing external stimulus (card value), while in the Necker Cube surprise emerges from an internal change to a constant external stimulus. The distinction between internal and external attribution of uncertainty (Kahneman & Tversky, 1982) is by-passed with the use of both a computational framework and neuroimaging. Cross-domain studies of decision-making are rare (Baasten et al., 2010) but we ventured across the sub-disciplines of neuroeconomics and psychophysics, motivated by two assumptions: first, the historical Helmholtzian view of inferential processes in relation to brain function, and second, the view that computational accounts of decision-making, if true, should hold for any decision type (O’Connell et al., 2018).

### Perceptual Switches as Involuntary Decisions

Several fMRI studies have probed perceptual switches (Frässle et al., 2014; Sterzer et al., 2009), but we examine their underlying, latent decision variables with a computational framework applied to fMRI signals, which allows us to empirically capture the hidden process of inference. The Necker Cube provokes sharp transitions between perceptual states, providing an ideal stimulus to capture perceptual error, unlike binocular rivalry, where more gradual transitions exist, introducing a temporal ambiguity in relation to the emergence of the error (e.g., Leopold et al., 1998; Naber et al., 2011). While perceptual switches are thought to be spontaneous and involuntary (Sterzer & Kleinschmidt, 2007), there are questions on whether reversals are subject to volitional control (Hugrass & Cuther, 2012; van Ee et al., 2005). In the study above, participants are encouraged to view stimuli passively, to control for willful perceptual switches. Further, dominance times recorded follow a stochastic time-course, suggesting spontaneous switches in perception.

### Perceptual Switches as Prediction Errors

By casting perceptual switches as prediction errors, we allay controversies regarding top-down (Wang et al., 2013; Long & Toppino, 2004; Sterzer et al., 2009) versus bottom-up (Polonsky et al., 2000; Parkonen et al., 2008; Pearson et al., 2007) processing in ambiguous stimuli, as the iterative process of prediction and error exchanges information between low- and high-level areas (Rao & Ballard, 1998; Summerfield & de Lange, 2014). Nonetheless, few studies have explicitly modeled switches as prediction errors. Sundareswara and Schrater (2008) characterize switches as inferential processes by applying a Markov Renewal Process to Necker Cube dominance times. In an fMRI study, a Bayesian account modeled switches in Lissajous figures as prediction errors (Weilnhammer et al., 2017). While both stimulus and formal account differed from ours, Weilnhammer and colleagues nonetheless find a response in the anterior insula to perceptual switches, in line with our results.

### Parsimony in Model Choice

The brain’s role in decision-making demands a parsimonious computation of uncertainty. Various models have been used to quantify uncertainty (Rao, 2010; Kepecs & Mainen, 2012; Vilares et al., 2012) but we opted to couch the mean-variance theorem (accounting for risk), within a predictive coding framework (to measure surprise), first to build on our previous work (Preuschoff et al., 2006; Preuschoff et al., 2008a; Preuschoff et al., 2011) but also due to its simplicity. In economics, the mean-variance framework encapsulates uncertainty by integrating both the mean and variance (risk) of a utility distribution (Kroll et al., 1984; Markowitz, 1952). From a homeostatic perspective, this framework presents an ideal model of uncertainty because decision-making with only two values leaves an agent neurally unencumbered, and accounting for risk facilitates learning, as estimating an option’s range shortens the trial-and-error process (D’Acremont & Bossaerts, 2008). It is worthwhile to note that several models can capture uncertainty (Friston, 2010) and some may be better suited to alternative paradigms, such as ones with an explicit learning component.

### Neural Specificity of First Order Prediction Errors

We replicate previous neuroimaging results for reward prediction error, notably its striatal correlate. Perceptual errors however correlate with a distinct response pattern in visual areas, suggesting these first-order errors elicit feature-specific neural responses; these errors nonetheless also implicate the hippocampus and temporal lobe, suggesting they recruit activity at the cognitive level, as would be expected in predictive coding (Clark, 2013). In contrast to reinforcement learning studies (Schultz et al., 2010; Kuhnen & Knutson, 2005; Carlson et al., 2011), we do not find a striatal involvement for perceptual error. This absence of striatal response in the perceptual domain supports a neural specificity to first-order prediction errors (Clark, 2013). The striatum has been hypothesized to act as a learning center and not just a reward hub (Balleine et al., 2007; Hare et al., 2008; Tricomi et al., 2009; Daniel & Pollman, 2014), an assumption that would see striatal responses to any prediction errors, including perceptual ones. Our results challenge this postulated role for the region. First-order prediction errors may account for case-specific decision features: BOLD responses in visual areas to perceptual prediction errors may reflect upstream, domain-specific, low-level errors (Parkkonen et al., 2008), while the insula may act as a common, downstream hub in the inference process.

### The insula in perceptual errors

Our model-based analyses on perceptual risk and surprise implicate the anterior insula in uncertainty, in line with previous studies on value-based decision-making (Preuschoff et al., 2006; Payzan-Le Nestour et al., 2013; Platt & Huettel, 2008) and perception (Sterzer & Kleinschmidt, 2010). Several studies investigating perceptual decision-making find increased reaction time correlated with the anterior insula, which corroborates the hypothesis that decision uncertainty relates to the region specifically (Ho et al., 2009; Binder et al., 2004; Thielscher & Pessoa, 2010). Sterzer & Kleinschmidt’s review on insular involvement in perceptual switches, titled “often observed but barely understood”, suggests salience elicits a consistent response in the region but bypasses its plausible role in inferential processes (Singer et al., 2009). We propose that the insula responds to ambiguous perception because it is tuned to a conscious uncertainty. This insight is important because a formal framework can allow us to test populations that are less susceptible to illusions, as in autism (Pellicano & Burr, 2012) and schizophrenia (Schmack et al., 2015).

### The insula in inference

A considerable body of evidence implicates insular BOLD responses in a wide range of functions including language (Ackermann & Riecker, 2010; Ardila et al., 2014); auditory processing (Bamiou et al., 2003); pain (Peyron et al., 2000; Corradi-Dell’Acqua et al., 2011); disgust (Wicker et al., 2003); gustatory function (Small, 2010); perception (Sterzer & Kleinschmidt, 2010); decision-making (Weller et al., 2009; Preuschoff et al., 2008; Singer et al, 2009; Volz et al., 2005); uncertainty (Critchley et al., 2004; Xue et al., 2010; Jones et al., 2011; Weller et al., 2009); and emotion in general (Gasquoine, 2014). To account for all these functions, the insula has been cast as an interoceptive center, integrating bodily states into awareness (Craig & Craig, 2009). Interoception presents a specific, if broad, category that readily explains salient emotions (love, fear) as well as sensory states (gustation, pain). Awareness of said states, which generally includes a declarative component, may explain the insula’s function in language processing. But then what of the insula’s contribution to uncertainty, prediction errors and perception (Klein et al., 2013)? While error is generically viewed as a conflict or mistake, from a computational perspective it is simply a difference between two states. In a related manner, the insula constitutes a main component of the salience network in resting state fMRI (Menon & Uddin, 2010). Salience implies, at minimum, a deviation from a neutral state. We hypothesize that the insula’s involvement in uncertainty reflects its role in mediating between upstream prediction errors (irrespective of origin) and declarative states. Thus, if an agent can label a state and the latter arises from a prediction error, we would expect insular involvement. A notable feature of the insula is that it likely does not have an animal homologue (Craig, 2009, but see Nieuwenhuys, 2012), underscoring its role in high-level functions, including consciousness (Craig & Craig, 2009; Seth, 2013).

### Inference and consciousness

The spectre of Helmholtzian views on inference hovers over our principal question: does the brain infer reality in a general manner irrespective of stimulus type (visual, monetary) or end goal (adequate perception, or monetary gain)? The inferential process should be the cornerstone of our interaction with the environment, regardless of domain. Our aim was to exploit a parsimonious economic model of uncertainty in perceptual and financial decision-making to probe the inferential process assumed in both. Our perceptual task eschews key aspects of economic decisions: it is unconscious, involuntary and quasi-immediate. Nonetheless, the key element of uncertainty allows us to quantify the inferential process and extract measures of risk, as is commonly done in economic contexts. Crucially, perceptual risk does not remain a mere theoretical quantity: we edify our hypothesis with an insular BOLD response, mimicking the neural correlates of (economic) risk processing in previous studies. In examining the insula’s functional role, we hypothesize that it is uniquely responsible for *conscious inference*, the result of an inference that can be recognized, as with a perceptual switch or the outcome of a gamble. Our results overall suggest that uncertainty can be quantified with a common framework across functional domains and further implicates a shared neural system, supporting the theoretical notion that the brain acts as an inference machine.

## Acknowledgments

This work was supported by the Swiss National Science Foundation (135687) and the German Research Foundation (DFG; grant no.: EI 852/3-1).

