## SupplementaryInformation for "Anterior Insula Reflects Inferential Errors in Value-Based Decision-making and Perception"

L. Loued-Khenissi et al., 2019

A.1

In the Necker Cube continuous task, perceptual prediction errors, as well as perceptual risk prediction errors were included as parametric modulators of switch response event onsets to highlight each prediction errors respective contribution to the BOLD response at that time. At the same event onset, we test for the 0 order effect of switch response with a one-sample t-test for that regressor. Below are results of that test, thresholded at p = 0.05 FWE.

| Contrast Switch (p = 0.05, FWE) | | | | | | | | | | |
| --- | --- | --- | --- | --- | --- | --- | --- | --- | --- | --- |
| cluster | cluster | cluster | peak | peak | peak | peak |  |  |  |  |
| p(FWE-corr) | equivk | p(unc) | p(FWE-corr) | T | equivZ | p(unc) | x | y | z {mm} |  |
| 0 | 302 | 0 | 0.001 | 9.1 | 5.66 | 0 | -30 | -54 | -28 | L Cerebellum Exterior |
|  |  |  | 0.001 | 8.66 | 5.52 | 0 | -40 | -62 | -28 |  |
|  |  |  | 0.002 | 8.51 | 5.47 | 0 | -20 | -60 | -18 |  |
| 0 | 268 | 0 | 0.001 | 9.06 | 5.65 | 0 | -44 | 16 | -4 | L Frontal Operculum |
|  |  |  | 0.009 | 7.47 | 5.11 | 0 | -54 | 4 | 4 |  |
|  |  |  | 0.031 | 6.7 | 4.8 | 0 | -48 | -2 | 10 |  |
| 0 | 118 | 0 | 0.002 | 8.43 | 5.44 | 0 | 32 | -54 | -32 | R Cerebellum Exterior |
|  |  |  | 0.004 | 8.03 | 5.31 | 0 | 44 | -58 | -30 |  |
|  |  |  | 0.022 | 6.91 | 4.88 | 0 | 24 | -58 | -22 |  |
| 0 | 72 | 0.001 | 0.005 | 7.82 | 5.24 | 0 | 48 | -40 | 32 | R Supramarginal Gyrus |
|  |  |  | 0.046 | 6.46 | 4.7 | 0 | 54 | -38 | 22 |  |
| 0 | 78 | 0.001 | 0.011 | 7.32 | 5.05 | 0 | 50 | 12 | 0 | R Frontal Operculum |
|  |  |  | 0.023 | 6.9 | 4.88 | 0 | 58 | 10 | 4 |  |
| 0.001 | 34 | 0.019 | 0.023 | 6.88 | 4.87 | 0 | -10 | 20 | 36 | L Supplementary Motor Cortex |

Table A.1 Whole-brain results of one-sample t-test performed on perceptual switch onsets (p=0.05, FWE corrected)

A.2 Report vs Replay

The perceptual switch as reported includes a motor response; our modulation of the switch is done to capture its degree of uncertainty. Thus we measured exogenous switch related BOLD responses by introducing replay trials as control conditions. Replay trials present disambiguated versions of the Necker Cube, presented at the same times as previously reported endogenous switch events. The replay condition is meant to capture BOLD responses related to viewing the cubes, deciding to report on a cube’s conformation and related reaction times. Contrasts for report versus replay conditions show no significant activation for report > replay, but do yield widespread bilateral activations in angular, frontal and middle temporal gyrii in the replay > report condition. Our results for this contrast are, interestingly, in line with previous findings that suggest perceptual ambiguity dampens neuronal spiking activity (Emadi & Esteky, 2013). We nonetheless account for replay responses by including them as regressors in our analyses.

| cluster | cluster | cluster | peak | peak | peak |  |  | null | Region |
| --- | --- | --- | --- | --- | --- | --- | --- | --- | --- |
| p(FWE-corr) | equivk | p(unc) | p(FWE-corr) | T | p(unc) | x | y | z {mm} |  |
| 0 | 969 | 0 | 0.001 | 10.14 | 0 | -54 | -56 | 42 | Angular  Gyrus L |
|  |  |  | 0.028 | 7.3 | 0 | -42 | -58 | 40 |  |
| 0 | 1850 | 0 | 0.002 | 9.24 | 0 | 46 | 30 | 30 | Mid Frontal Gyrus R |
| 0 | 871 | 0 | 0.041 | 7.04 | 0 | -2 | 32 | 46 | Superior Frontal Gyrus |
| 0 | 1100 | 0 | 0.045 | 6.98 | 0 | -46 | 12 | 44 | Mid Frontal Gyrus L |
| 0 | 1290 | 0 | 0.055 | 6.84 | 0 | 36 | -52 | 40 | Angular  Gyrus R |
| 0.001 | 518 | 0 | 0.327 | 5.59 | 0 | 60 | -34 | -6 | MTG R |
| 0 | 594 | 0 | 0.518 | 5.21 | 0 | 36 | -80 | -10 | Occ. Fusiform Gyrus R |
| 0.038 | 238 | 0.004 | 0.638 | 5 | 0 | -58 | -32 | -10 | MTG L |

Table A.2 Whole Brain Analyses for Replay > Report condition ( cluster corrected, with a starting threhsold of p =.001, k = 25)

A.3 Dominance Time Distributions

Necker Cube dominance times vary within and across individuals and generally follow a log-normal or gamma distribution. Below, we fit dominance times across all subjects, separated by stimulus ambiguity level (20%/80%; 35%/65%; 50%) to gamma distributions.

Figure A.3 Dominance Time Distributions Across Differing Stimulus Ambiguities. Dominance times for all participants were pooled and binned according to stimulus type (least, partial and most ambiguous). These values were then fit to a gamma distribution to ensure that the Necker Cube manipulation yielded dominance times that conform to expected, spontaneous switches.

| MLE (Gamma Distribution) | | |
| --- | --- | --- |
|  | k | µ |
| All Dwell | Times 1.5457 | 3.2922 |
| Ambiguous Stimulus Dwell Times (50%) | 1.5975 | 2.8688 |
| Partially Ambiguous Stimulus Dwell Times (35%/65%) | 1.5943 | 3.1817 |
| Disambiguated Stimulus Dwell Times (20%/80%) | 1.4834 | 3.7866 |

Table A.3 –Maximum likelihood estimates for gamma distribution parameters of dwell times

across subjects in all stimulus conditions as well as ambiguous, partially ambiguous and

disambiguated stimulus conditions.

A.4

Alternative Models of Surprise and Risk

Models of surprise as probabilistic expectation violations include unsigned prediction error and Shannon information, in addition to risk prediction error. In our study, these different models are highly correlated, as shown in the Figure A.4.1 below. Similarly, risk as variance in our study is highly correlated with Shannon entropy (Figure A.4.2), limiting the need for model comparison.

Equation A.4.1

$$Is= -log(p\left( outcome | Card1, Bet \right))$$

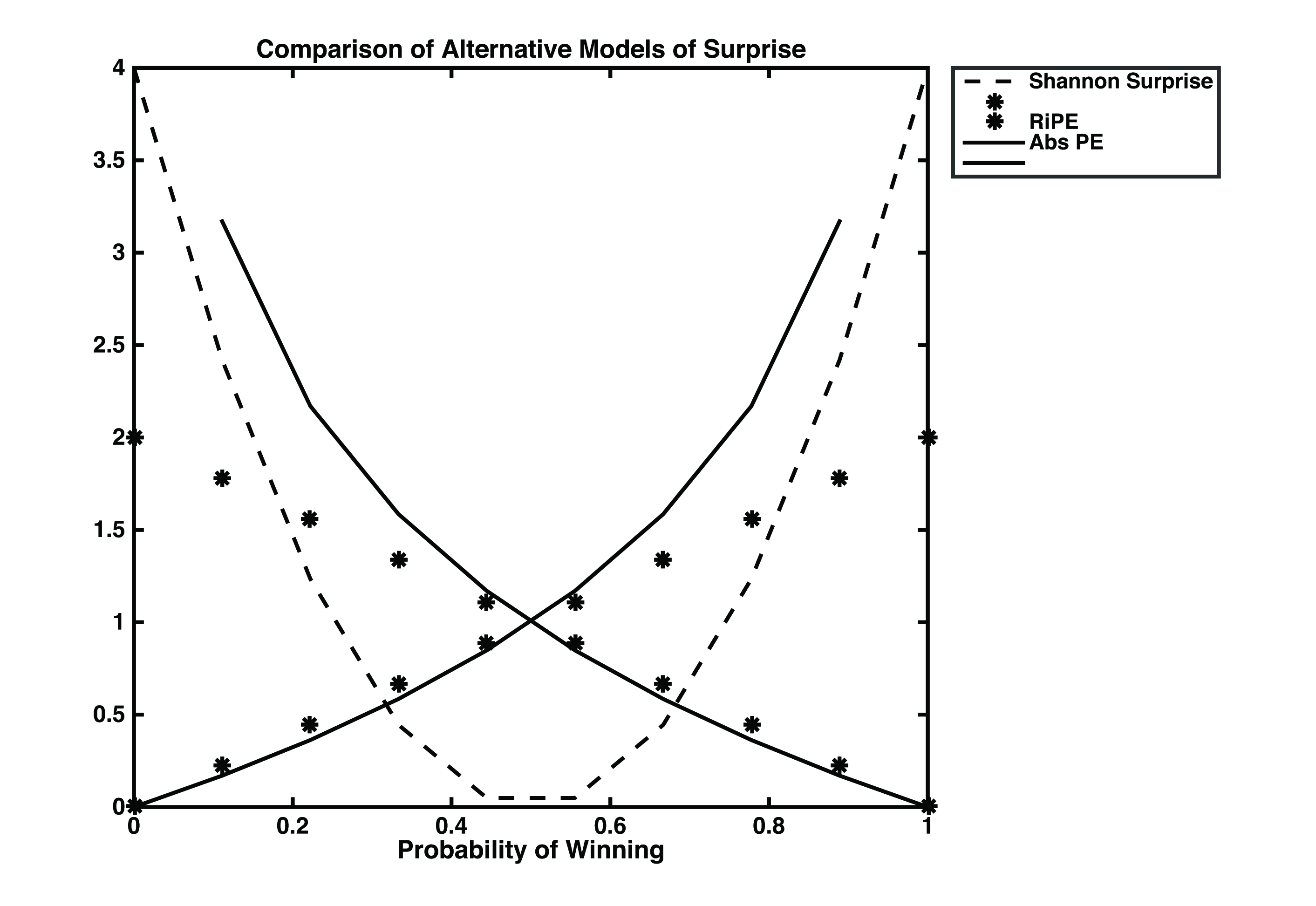

Figure A.4.1. Shannon Surprise, risk prediction error and absolute prediction error across probabilities of winning for all possible win and loss outcomes.

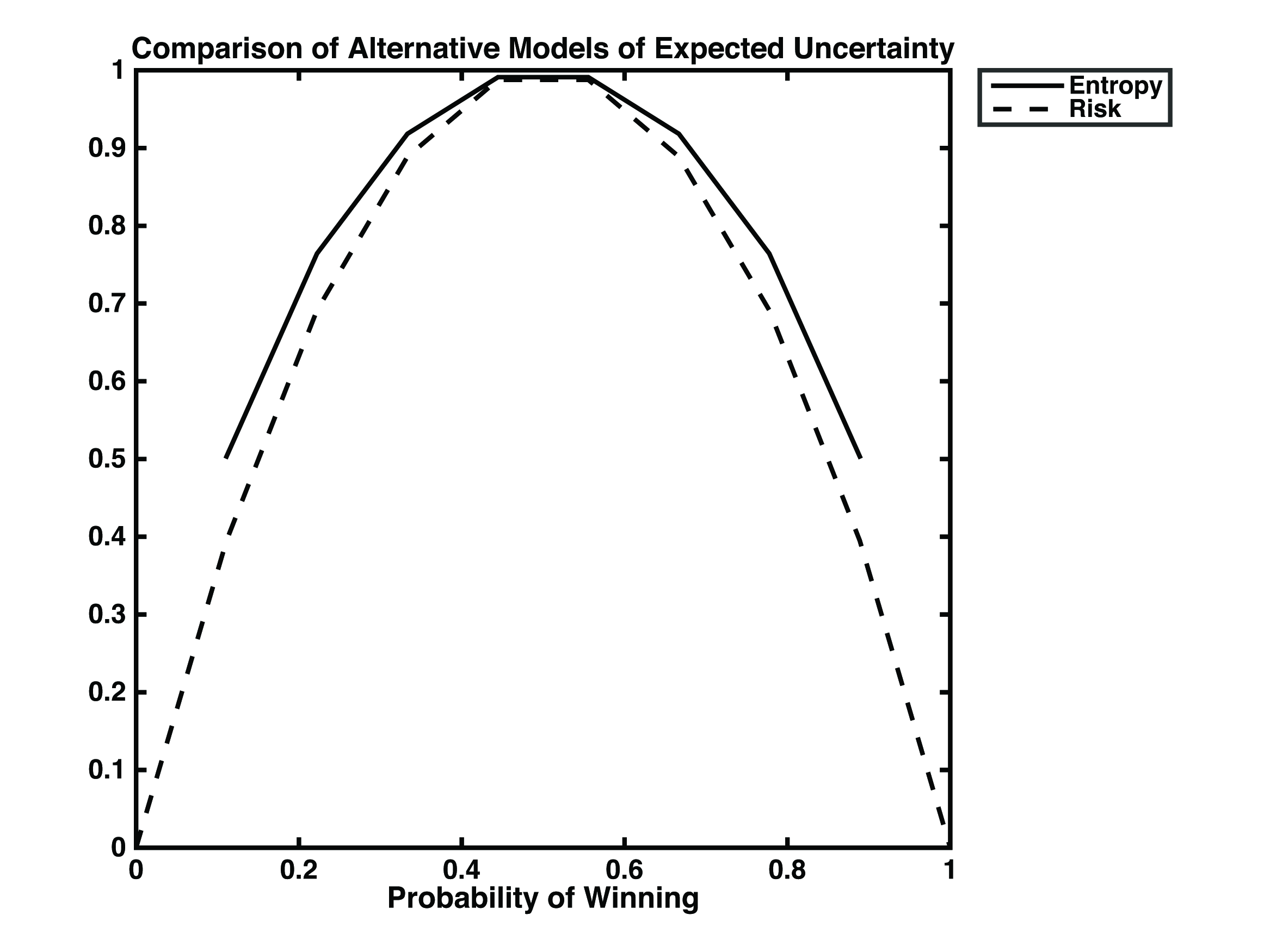

Figure A.4.2 Risk and entropy plotted according to the range of probabilities of winning in the Card Game. While the two values nearly overlap, it is worthwhile to not that certain trials (when Card 1 is 1 or 10) yield undefined values of entropy.

Equation A.4.2

$$H= -p_{win}*l\mathrm{og}_{2}(p_{win}) - (1 -p_{win})*l\mathrm{og}_{2}(1 - p_{win})$$
